# Interplay between medial nuclear stalling and lateral cellular flow underlies cochlear duct spiral morphogenesis

**DOI:** 10.1101/2020.07.24.219469

**Authors:** Mamoru Ishii, Tomoko Tateya, Michiyuki Matsuda, Tsuyoshi Hirashima

**Affiliations:** Laboratory of Bioimaging and Cell Signaling, Graduate School of Biostudies, Kyoto University; Department of Speech and Hearing Sciences and Disorders, Faculty of Health and Medical Sciences, Kyoto University of Advanced Science; Department of Pathology and Biology of Diseases, Graduate School of Medicine, Kyoto University; Japan Science and Technology Agency, PRESTO

**Keywords:** Differential growth, FRET imaging, Interkinetic nuclear migration, MAPK/ERK, Mathematical model, Morphogenesis, Spiral pattern formation

## Abstract

A notable example of spiral architecture in organs is the mammalian cochlear duct, where the duct morphology is critical for hearing function. Molecular genetics has revealed the necessary signaling molecules for the formation of spirals in organs, but it remains unclear how cellular dynamics generate bending and coiling of the cochlear duct during development. Here we show two modes of multicellular dynamics underlying the morphogenetic process by combining deep tissue live-cell imaging, Förster resonance energy transfer (FRET)-based quantitation, and mathematical modeling. First, surgical separation of the cochlear duct revealed that bending forces reside primarily in the medial side of the duct. In the medial pseudostratified epithelium, we found that nuclei stall at the luminal side during interkinetic nuclear migration, which would cause the extension of the luminal side, thereby bending the duct. Second, long-term organ-scale FRET imaging of extracellular signal-regulated kinase (ERK) activity showed that helical ERK activation waves propagate from the duct tip concomitant with the reverse multicellular flow in the lateral side of the duct, resulting in advection-based duct elongation. We propose an interplay of distinct multicellular behaviors underpinning spiral morphogenesis in the developing cochlear duct.

## Introduction

Spiral shapes are a widely occurring motif in many varied biological tissues and organisms, including shells, horns, and plants, but how spiral shapes form has remained unclear (Thompson, 1942). The general principle of spiral formation is differences in growth rate between the outer and inner tissue of the extending organ, with the growth rate of the outer tissue being faster than that of the inner one, which has been theoretically and experimentally demonstrated in shells and plants (Johnson et al., 2019; Raup and Michelson, 1965; Smyth, 2016; Wada and Matsumoto, 2018). The cellular processes causing this differential tissue growth are different in each organ (Johnson et al., 2019; Saffer et al., 2017), and so identifying the organ-specific mechanisms underlying these differential tissue growth rates is crucial to understanding the developmental process of spiral morphogenesis.

An example of a spiral organ is the mammalian cochlear duct, which is a tonotopically organized auditory organ in the inner ear (Fig. 1A). During murine development, the cochlear duct, composed of epithelial cells, elongates, bends, and coils to form a spiral. The molecular basis for the morphogenesis of the cochlear duct has been the subject of several previous studies. Gene knockout studies have clarified that the elongation of the cochlear duct requires sonic hedgehog (SHH) signaling from the cochleovestibular ganglion in the conical central axis of the cochlea (Bok et al., 2013; Liu et al., 2010; Tateya et al., 2013), fibroblast growth factor (FGF) signaling of epithelial cells (Pauley et al., 2003; Pirvola et al., 2000; Urness et al., 2018, 2015), and non-canonical Wnt–planar cell polarity (PCP) signaling of prosensory cells (Mao et al., 2011; Montcouquiol and Kelley, 2020; Qian et al., 2007; Saburi et al., 2008; Wang et al., 2005). Deletion of *Shh* expression leads to a shortening of the cochlear duct and a significant decrease in cell proliferation exclusively at the base region (Bok et al., 2013). In *Fgf10* null mutant mice, the cochlear duct is remarkably shorter, but cell proliferation is unaffected (Urness et al., 2015), suggesting that cell proliferation and other cellular processes regulate ductal outgrowth. From embryonic day (E) 14.5 onwards, the mediolateral active migration of prosensory cells, during which these cells intercalate radially with their neighbors (known as convergent extension), contributes to longitudinal duct extension in a Wnt–PCP pathway-dependent manner (Chen et al., 2002; Cohen et al., 2019; Driver et al., 2017; Yamamoto et al., 2009). This cell intercalation drives ductal elongation; however, it cannot explain the asymmetrical morphogenetic mode underlying the duct bending before E14.5. Although the underlying signaling pathways are well characterized, the physical cellular mechanisms underlying spiral morphogenesis of the cochlear duct remain elusive. In the present study, we aimed to identify the multicellular dynamics giving rise to the bending and coiling of the developing cochlear duct using a combination of live cell-imaging, FRET quantitation, and mathematical modeling.

**Figure 1.**
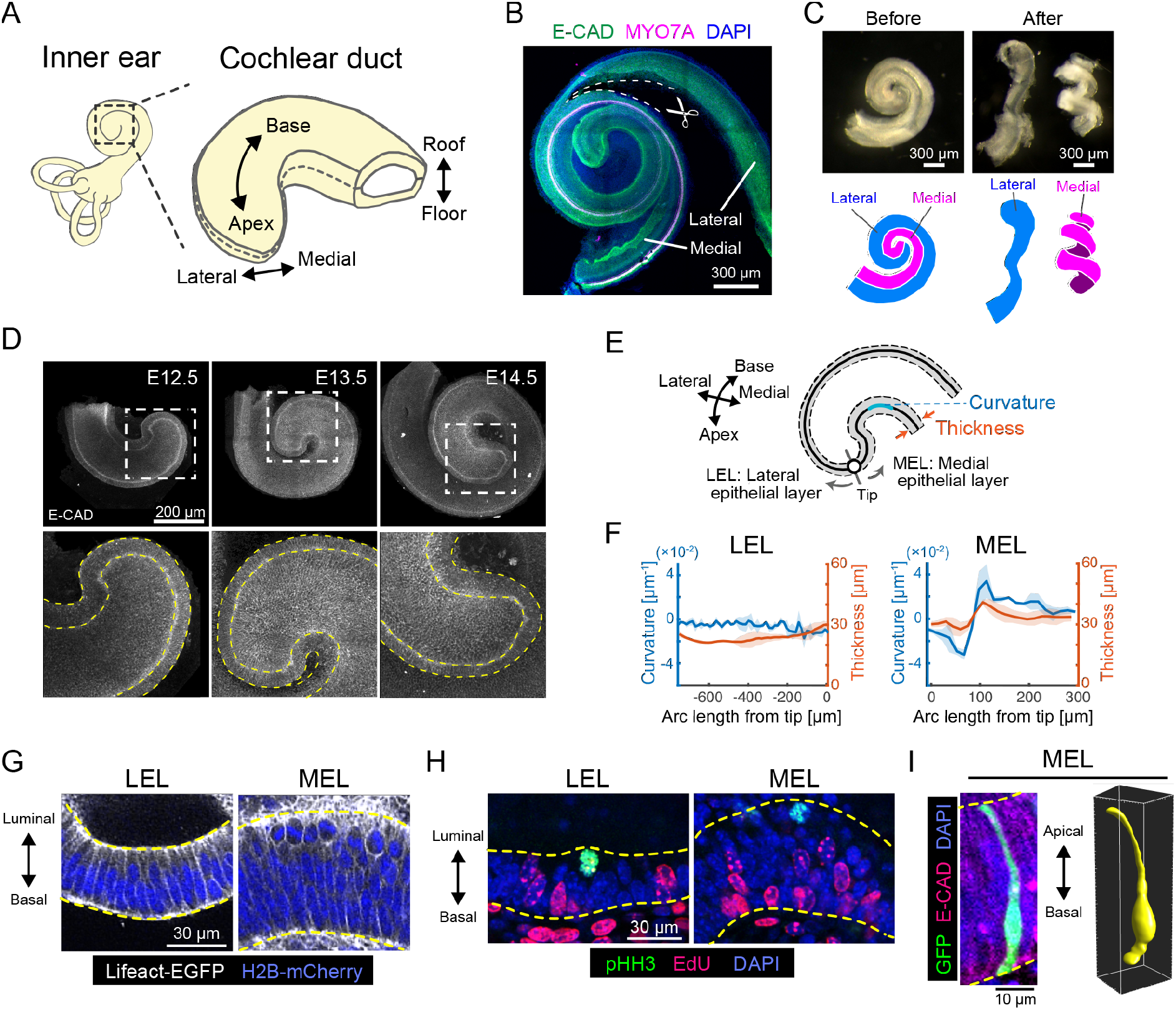
The pseudostratified medial epithelial layer curves in the cochlear duct. (A) Schematic diagrams showing the tissue axis and labels of the cochlear duct. (B) Immunofluorescence image of anti-E-cadherin (green) and anti-Myosin VIIA, a marker for sensory hair cells (magenta), with nuclear counterstaining using DAPI (blue) in the cochlea at E17.5 during tissue separation, with cuts represented by dashed lines. N=3 was confirmed. Scale bar, 300 μm. (C) Images of the cochlear duct before and after tissue separation. Stereomicroscope images (upper) and the corresponding illustrations of the lateral (blue, left) and medial (magenta, right) layers. N=3 was confirmed. Scale bar, 300 μm. (D) Immunofluorescence images of anti-E-cadherin staining in the murine developing cochlea. The lower rows are magnified images of the dotted squares in the upper rows. Yellow dotted lines represent the edges of the epithelial layer. Scale bar, 200 μm. (E) Schematic diagram showing regions used for morphological quantification. (F) Curvature and thickness as a function of the arc length from the tip along the lateral epithelial layer (LEL, left) and the medial epithelial layer (MEL, right) at E12.5. Mean ± standard deviation (s.d.) N=3. (G) Images of Lifeact-EGFP (white) and H2B-mCherry (blue) showing cell shape and nuclear position in the LEL and MEL. Yellow dotted lines represent the luminal and basal edges. Scale bar, 30 μm. (H) Fluorescence labeling images of anti-pHH3 (green) and EdU (magenta) with nuclear counterstaining using DAPI (blue) in the LEL and MEL. Yellow dotted lines represent the luminal and basal edges. Scale bar, 30 μm. (I) Fluorescence image of GFP transfected cells labelling single cells (green) and anti-E-cadherin (magenta) with nuclear counterstaining using DAPI (blue) in the MEL (left) and the corresponding 3D rendered image (right). Yellow dotted lines on the left represent the luminal and basal edges. Scale bar, 10 μm.

## Results

### The medial epithelial layer causes cochlear duct bending

Mediolateral asymmetry characterizes the spiral form of the cochlear duct (Fig. 1A). We therefore first examined the force balance between the medial and the lateral tissue by surgically separating the cochlear duct along the lateral side of the hair cells (Fig. 1B); the medial side included Kölliker’s organ and the prosensory domain, and the lateral side included the outer sulcus after separation. This manipulation led to the release of the mechanical stress affected mutually between the medial and the lateral tissue, enabling us to inspect the internal stresses of local tissues. The separated medial tissue curled much more after separation than before, while the lateral tissue was relatively uncurled (Fig. 1C and Mov. 1), indicating that active bending force are applied from the medial side, but not from the lateral side.

We next examined the morphology of the developing cochlear duct from E12.5 to E14.5 by staining an epithelial marker, E-cadherin, and performing organ-scale 3D imaging. During this developmental period, the cochlear duct elongates and coils without changes to the mediolateral width (Fig. 1D). The curvature and thickness of the epithelial layer are significantly larger on the medial side than on the lateral side in a horizontal section of the roof-floor axis (Fig. 1D). Hereafter, we refer to the epithelial layers on the medial and lateral sides as the medial and lateral epithelial layers (MEL and LEL), respectively (Fig. 1E). Morphological quantification revealed that the curvature and thickness of the MEL are highest at a point more than 100 μm away from the tip of the duct along the MEL (Fig. 1F and S1). In contrast, the curvature and thickness of the LEL remain constant (Fig. 1F and S1). These observations prompted us to explore the structure and dynamics of the MEL at single cell resolution.

Zooming in the epithelial layer clarified that most of the nuclei in the LEL align within a few nucleus diameters of each other (~20 μm) (Fig. 1G, left). However, nuclei in the MEL distribute broadly across a ~50 μm range, and mitotic cell rounding only occurs on the luminal side of the MEL (Fig. 1G, right). We also found that nuclei that stained positive for the M phase marker phospho-histone H3 (pHH3) were localized only on the luminal side of the epithelial layers, while nuclei in the S phase cells labeled with a short pulse of ethynyl deoxyuridine (EdU) were distributed predominantly on the basal side (Fig. 1H). Mosaic cell labeling revealed that single cell bridges extended between the luminal and basal edges of the MEL via long protrusions (Fig. 1I), clearly indicating that the MEL is a proliferative pseudostratified epithelial tissue.

### Luminal nuclear stalling promotes MEL bending

We next focused on the cellular dynamics in the MEL with a two-photon microscope. For live imaging of the MEL, the outer cartilaginous shell that will develop into the bony labyrinth was completely removed to expose the cochlear duct prior to *ex vivo* culture. We used a reporter mouse line ubiquitously expressing fluorescence proteins localized in the cytoplasm, allowing us to recognize the nuclei as the regions without fluorescence, as well as the luminal and basal edges of MEL in the pseudostratified epithelium. We observed differences in the nuclear movement between the flat region proximal to the tip of the apex and the curved region distal to the tip of apex (Fig. 2A, Mov. 2). The nuclei moved to the luminal edge of the epithelial layer before cell division, and after cell division the daughter nuclei returned to the basal side in the flat region (Fig. 2B). In the curved region, however, the cell nuclei remained at the luminal side even after cell division (Fig. 2C). To make this difference clearer, we tracked the nuclei and quantified their distance from the luminal edge at 3 hours after nuclear division. After classifying the tissue into two categories – flat and curved – based on the half-maximal curvature (Fig. 2D), fewer nuclei move to the basal side in the curved region than in the flat region (Fig. 2D’), suggesting that there are two different modes of luminal-basal nuclear movement in the MEL, depending on the distance from the duct tip. In addition, we examined two angles of cell division orientation, φ and θ, in the 3D polar coordinates from the live imaging data, with φ and θ representing the zenith and azimuth angle, respectively (Fig. 2E). The φ angle was between 0° to 45°, indicating that cell division mostly occurs in parallel to the luminal surface of the MEL (Fig. 2F). The distribution of θ shows that most cells divide along the apex-base axis rather than along the roof-floor axis (Fig. 2G), suggesting that the orientation of cell division contributes to the apex-base local expansion of the luminal side of the MEL.

**Figure 2.**
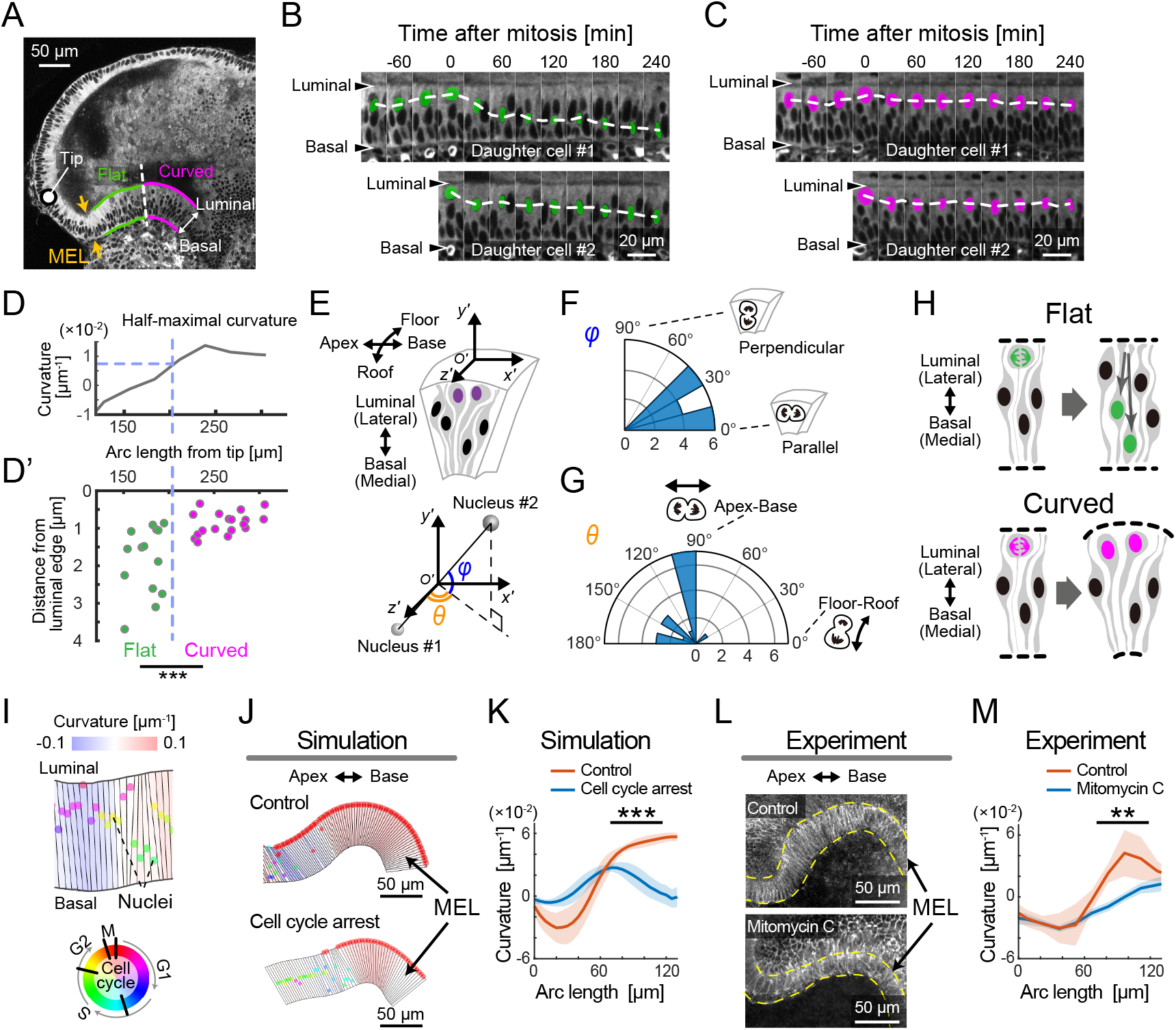
Live imaging and mathematical modelling demonstrate that luminal nuclear stalling promotes MEL bending via luminal-basal differential growth. (A) Image of a section of an E14.5 cochlea in cytoplasmic reporter mice (CFP channel of ERK-FERT mice). The MEL was classified into flat (green) and curved (magenta) regions and the boundary between the flat and curved regions is indicated by a dotted line. The circle indicates the duct tip. Scale bar, 50 μm. (B, C) Kymographic images of the flat region (B) and the curved region (C). Dotted lines denote a change in position of manually marked nuclei over time. Scale bars, 20 μm. (D, D’) For 16 nuclear divisions that could be tracked, the distance from the luminal edge 3 hours after nuclear division was plotted over the arc length from the tip (D’) together with the MEL curvature (D). At half-maximal curvature (D), the samples in (D’) were classified into two groups, i.e., flat (n=14) and curved (n=18). Mann–Whitney U test, p=0.00038. (E) Nuclear division orientation is represented by two angles, φ and θ, in the polar coordinate O’. (F, G) Angle distribution of φ and θ. n=16. Rayleigh test, p=0.00096 for (F) and p=0.15 for (G). (H) Schematics for the expected morphology depending on the mode of nuclear movement. IKNM would lead to the flat (upper) and asymmetric IKNM would lead to the curved (lower). (I) Virtual epithelial layer in the model simulation. The colored circles denote nuclei with the cell cycle state represented in the bottom. (J, K) Model prediction in the cell cycle arrest before S phase entry in the growing virtual MEL. Representative images (J) and the curvature over the arc length (K). Mean ± s.d. N=10. Mann–Whitney U test, p<0.001. Scale bar, 50 μm. (L, M) The effect of mitomycin C treatment. Representative images of resultant cochlear duct immunostaining of anti-E-cadherin (J) and the curvature over the arc length (M). Mean ± s.d. N=3. Mann–Whitney U test, p=0.0017. Scale bar, 50 μm.

Luminal-basal nuclear movement in the flat region exhibits the behavior known as interkinetic nuclear migration (IKNM), which has been observed in various pseudostratified epithelia (Grosse et al., 2011; Kosodo et al., 2011; Meyer et al., 2011; Norden et al., 2009; Sauer, 1935). During this process, nuclei at the basal side move close to the luminal surface prior to mitosis, and the daughter nuclei move back to the basal side, which results in even growth within the epithelial layer (Fig. 2H, upper). This mode of nuclear migration would not necessarily result in bending of the epithelial tissue. In contrast, lack of basalward movement during IKNM, which we refer to as ‘luminal nuclear stalling’ hereafter, results in asymmetric one-way flux of nuclei from the basal side to the luminal surface of the layer, which results in local cell crowding and expansion due to the division occurring at the luminal side (Fig. 2H, lower). This asymmetrical IKNM, along with oriented cell division, may contribute to differential growth between the luminal and basal sides of the MEL, and thus cause the physical bending of the MEL.

To test this possibility, we built a cell-based mechanical model and examined whether nuclear movements affect the curvature of an epithelial layer. Single epithelial cells in the layer were represented as polygons with vertices consisting of the luminal and basal edges, which could deform according to the cell cycle-dependent nuclear position, based on mechanical interactions between neighboring cells (Fig. 2I). We then introduced a parameter ***γ*** controlling the degree of basalward movement after IKNM – the nucleus moves to the basal edge when ***γ*** = **1** and stays at the luminal side when ***γ*** = **0**. Our mathematical simulation demonstrated that the curvature of the epithelial layer monotonically decreases as a function of ***γ*** (Fig. S2A), which supports our observation that luminal nuclear stalling mechanically contributes to bending of the epithelial layer. We used an *in silico* experiment to predict that cell cycle arrest at the S-phase entry would decrease the curvature of the MEL (Fig. 2J, 2K, and Mov. 3). This prediction was validated by treating explant cochlea with the DNA synthesis inhibitor Mitomycin C. After 1 day of treatment with 10 μM Mitomycin C, EdU-positive S phase cells were no longer detected in the cochlea (Fig. S2B). Under these conditions, the amount of curvature along the MEL was significantly decreased in the curved region (Fig. 2L and 2M), which is consistent with the model prediction. These results corroborate our hypothesis that luminal nuclear stalling after luminal mitosis promotes MEL bending.

### Spatial heterogeneity of cell proliferation suggests cellular inflow to the lateral side of the apex to realize cochlear bending

One plausible mechanism for achieving the spiral morphology of the cochlear duct is MEL-driven differential growth, but another possibility is that cell proliferation might occur at a faster rate on the lateral side than on the medial side. To address this, we examined the spatial distribution of proliferating cells within 30 min of labeling of EdU on the roof and floor side of the cochlear duct (Fig. 3A and 3B). For quantification, the image domain of the cochlear duct was divided into interrogation regions and the averaged fluorescence intensity of labelled EdU was measured within each region (Fig. 3C). On the floor side, EdU-positive cells were more abundant in the medial side than in the lateral side around the apex (arrows, Fig. 3B and 3D, left). However, on the roof side of the cochlear duct, EdU-positive cells were distributed evenly, with slightly fewer cells at the base than at the apex and without a significant bias along the mediolateral axis (Fig. 3B and 3D, right). These observations negate the possibility that cell proliferation rates on the lateral side of the cochlear duct could drive mediolateral differential tissue growth and cause duct bending. The spatial map of EdU intensity shows that cell proliferation rates were higher in the floor-base region (arrowheads, Fig. 3B and 3D, left). Supposing that cell proliferation is the main driver of local tissue growth, the higher volumetric growth observed in the medial side than in the lateral side would contribute to the duct bending inward at the lateral side, which contradicts the innate cochlear morphogenesis. We thus hypothesized that the cells in the lateral side of the growing apex may be supplied by proliferation ‘hot spot’ in the basal region of the cochlear duct, which could resolve the observed mismatch between tissue growth rates in the medial and lateral sides of the duct.

**Figure 3.**
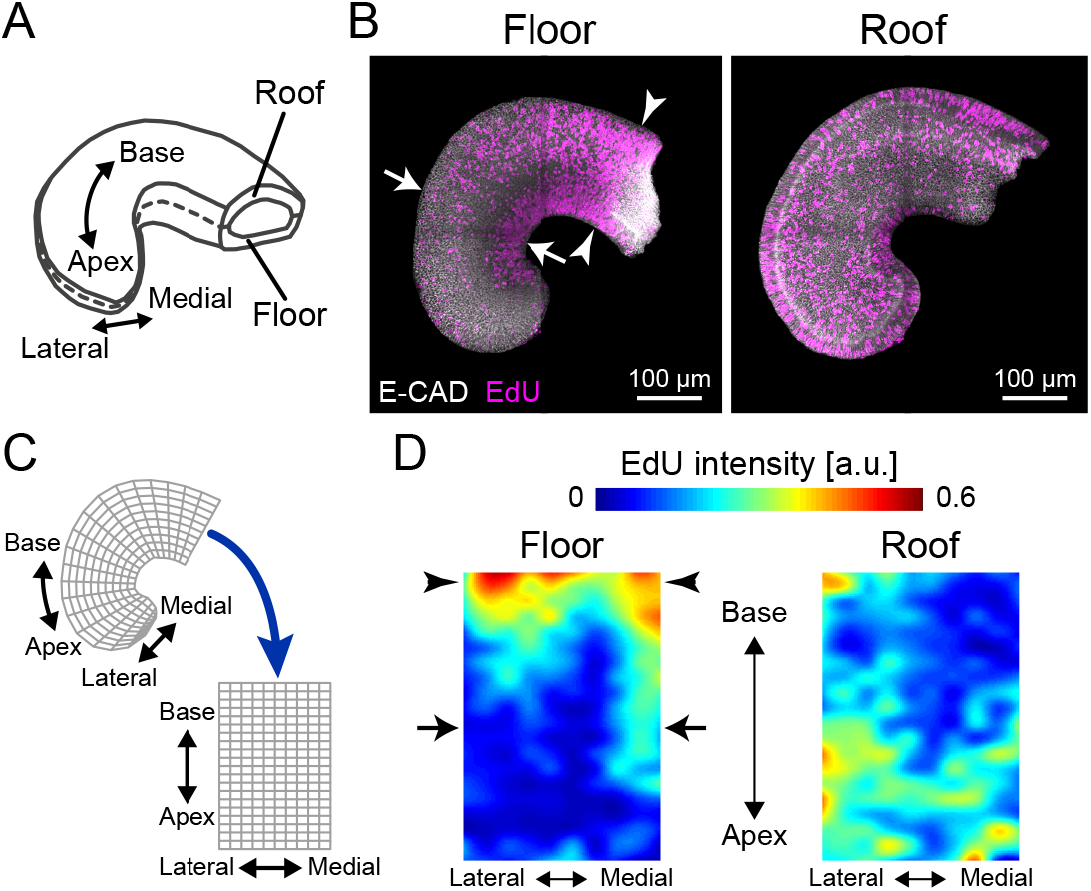
Cell proliferation mapping suggests cellular inflow to the lateral side of the cochlear duct apex to realize cochlear bending. (A) Schematic of tissue axis of the cochlear duct. (B) Maximum projection images of stained anti-E-cadherin (white) and EdU (magenta) in the roof and floor region of cochlear duct at E12.5. Representative images showing EdU signals in the cochlear duct are shown. Arrows indicate the region of EdU signal gradient from the medial to the lateral side of the duct. Arrowheads indicate the base region where the EdU intensity is concentrated. Scale bar, 100 μm. (C) Regional mapping from irregular grids in the cochlear duct to regular grids. (D) Heat map of EdU intensity in the roof and floor region of the duct. Arrows and arrowheads are the same as in (B). N=3 was confirmed.

### Retrograde helical ERK activation waves drive base-to-apex multicellular flow

How are cells supplied from the base of the cochlear duct to the lateral side? Earlier studies reported that FGF signaling is critical for cochlear duct outgrowth (Pirvola et al., 2000; Urness et al., 2015). We therefore focused on ERK MAP kinase, a downstream kinase in the FGF signaling pathway, and used a reporter mouse line that ubiquitously expresses a Förster resonance energy transfer (FRET)-based biosensor for ERK activity in the cytosol (Harvey et al., 2008; Komatsu et al., 2018, 2011). 3D FRET imaging using two-photon microscopy revealed that ERK is preferentially activated in the lateral-roof side of the cochlear duct, including the outer sulcus and stria vascularis (Fig. 4A, 4A’, and Mov. 4), which is consistent with the previously reported distribution of *Fgfr2* expression (Urness et al., 2015).

**Figure 4.**
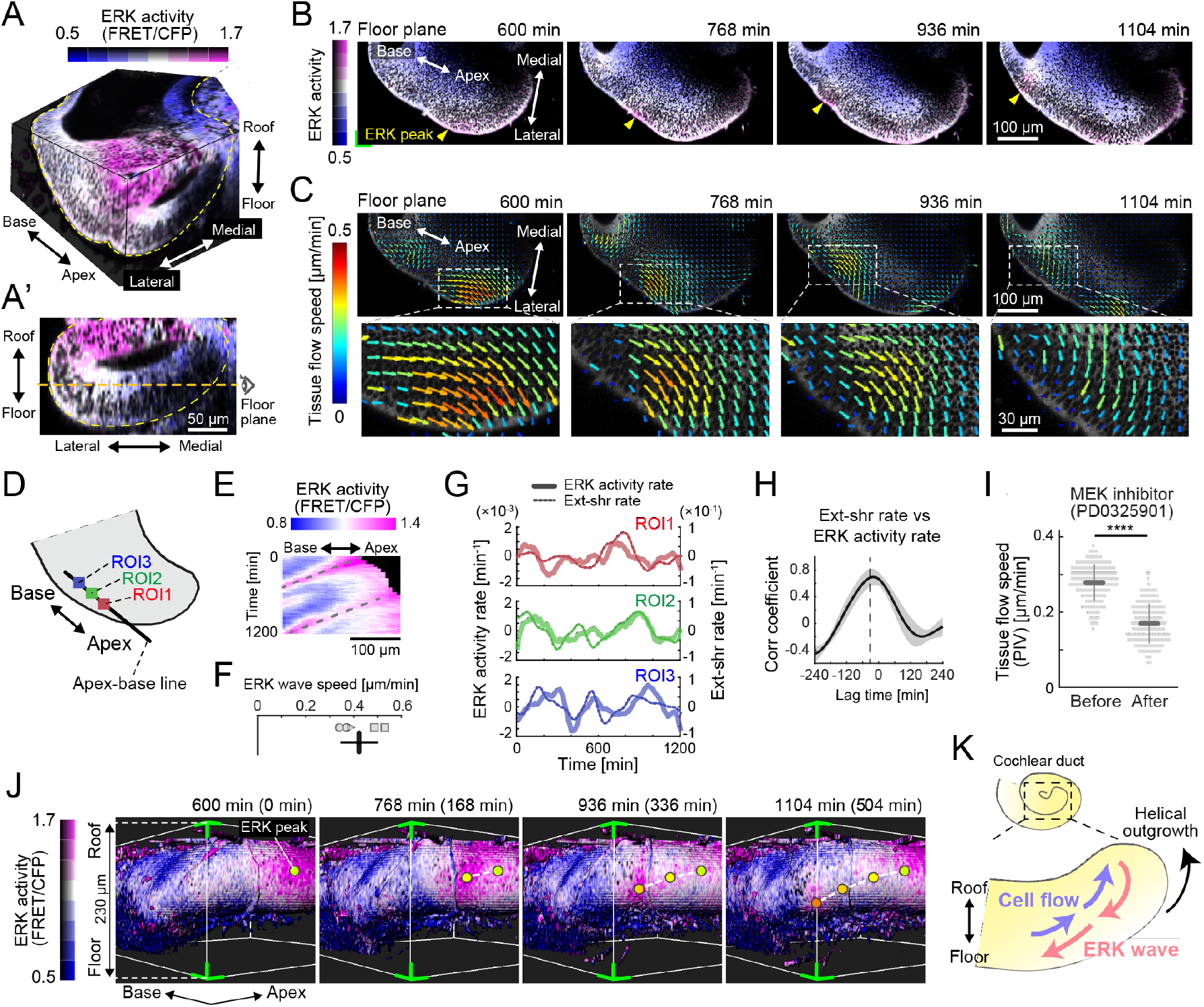
Retrograde helical ERK activation waves drive base-to-apex multicellular flow. (A) 3D ERK activity map in the cochlear duct at E12.5. (A’) Cross section view (medial-lateral and roof-floor plane) of (A). Orange dotted line indicates the floor plane shown in (B) and (C). Scale bar, 50 μm. (B) Time-lapse snapshots of ERK activity maps in the floor plane. Time indicates the elapsed time of live imaging. Yellow arrowheads indicate the ERK activity peak. Scale bar, 100 μm. (C) Time-lapse snapshots of tissue flow speed obtained by PIV in the floor plane. Scale bar, 100 μm. (D) Schematic diagram showing the axis, the apex-base line for kymography, and regions of interests (ROIs). (E) Representative kymograph of ERK activity. The horizontal axis indicates the position on the apex-base line shown in (D) and the vertical axis indicates the elapsed time of live imaging. Dotted lines represent oscillatory waves from the apex to the base. Scale bar, 100 μm. (F) ERK wave speed with mean and s.d. n=5 from N=3. (G) Time-series ERK activity rate and extension-shrinkage rate in representative 3 different ROIs. (H) Cross-correlation between the extension-shrinkage rate and ERK activity rate. n=12. Mean ± s.d. (I) Tissue flow speed before and after MEK inhibitor PD0325901 treatment at 1 μM. n=285 from N=3. Two-sample t-test, p<0.001. (J) Time-lapse snapshots of surface rendered ERK activity maps in the cochlear duct at E12.5. The green corners correspond to the green corner on the images shown in (B) and viewed from the left-bottom corner of (B). Circles indicate the position of ERK activity peaks and the connecting dotted lines indicate a trace of the peak shift. The time scale is the same as in (B). (K) Schematics for the ERK activity waves and cell flow.

For continuous observation during ductal outgrowth, we established an explant culture method in which the capsule above the duct tip was partially removed, allowing 3D organ-scale long-term imaging of ERK activity. Surprisingly, the time-lapse images of the cochlea dissected at E12.5 revealed that ERK activation propagates intercellularly as oscillatory waves from the apex to the base of the floor side (Fig. 4B and Mov. 5), while ERK is constitutively activated around the ductal tip of the roof side (Fig. S3A and S3B). We next quantified multicellular tissue flow by particle image velocimetry (PIV) at the supra-cellular (4–5 cell length) scale, and found that cells coherently move as clusters of ~100 μm diameter from the base to the apex of the floor side, again as oscillatory waves, and similarly to the ERK activation waves of the floor side (Fig. 4C and Mov. 5). In the roof side, the multicellular tissue flows directly toward the direction of elongation around the duct tip (Fig. S3C). Kymography of ERK activity along the apex-base line of the lateral-floor side (Fig. 4D) shows oscillatory retrograde ERK activity waves (Fig. 4E and Fig. S3D), which proceed at a speed of 0.42 ± 0.078 μm min^−1^ (mean ± standard deviation) in space-fixed coordinates (Fig. 4F).

Since ERK activation can be induced by cell extension during collective cell migration (Hino et al., 2020), we further calculated the ERK activity rate, i.e., the time derivative of ERK activity, and the extension-shrinkage rate, i.e., the local tissue strain rate along the apex-base line, by using time-series data of ERK activity and PIV speed, respectively (Fig. S3E). We found that both the ERK activity rate and the extension-shrinkage rate oscillate across the time course (Figure 4G). Moreover, cross-correlation analysis revealed that the local tissue deformation, represented by the extension-shrinkage rate, precedes ERK activity changes by 24 min on average (Fig. 4H). This is consistent with a regime of mechano-chemical coupling for collective cell migration: extension-triggered ERK activation promotes cell contraction, pulling the neighboring cells, which eventually evokes another round of ERK activation in the neighboring cells (Boocock et al., 2020; Hino et al., 2020). The observed lag time between the ERK activity rate and the extension-shrinkage rate can be reproduced using a minimal mathematical model of the mechano-chemical coupling, but not by an uncoupling model under which the ERK activation waves regulate the cell deformation unidirectionally, supporting the plausibility of mechano-chemical coupling rather than an uncoupling regime (Fig. S4). The role played by ERK was confirmed by treating the cochlear duct with an inhibitor of the upstream kinase MEK, PD0325901. Treatment with the MEK inhibitor at 1 μM resulted in a significant decrease of the tissue flow speed (Fig. 4I) as well as ERK inactivation (Fig. S3F), corroborating that ERK activation waves contribute to the base-to-apex multicellular tissue flow.

Finally, we extended the analysis to 3D dynamics of ERK activity and cell movement in the developing cochlear duct. Surface rendering of the ERK activity map in the cytosol indicated that ERK activity peaks shift from the apex-roof to the base-floor in the lateral side of the cochlear duct (Fig. 4J and Mov. 6). Concomitantly with helical ERK activity waves, coherent cell movements can be observed from the base-floor to the apex-roof in the opposite direction to the ERK waves (Mov. 6). This observation strongly suggests that ERK-mediated helical collective cell movement could drive 3D duct coiling underlying the spiral morphogenesis of the cochlear duct (Fig. 4K).

## Discussion

Previous genetic studies have revealed the molecular basis of cochlear duct elongation during development, but have been unable to explain the physical mechanisms by which the duct bends because of its severe phenotype (Bok et al., 2013; Groves and Fekete, 2012; Urness et al., 2018, 2015). Motivated by these earlier studies, we have visualized the cochlear duct development by two-photon microscopy and found that nuclei moving from the basal side stall at the luminal side after cell division, which gives rise to bending via differential tissue growth between the luminal and basal sides of MEL. We also elucidated that coherent cellular flow occurs from the base to the apex exclusively in the lateral wall of the growing cochlear duct, which is accompanied by retrograde ERK activation waves. Thus, our long-term deep tissue imaging has illuminated unprecedented dynamics of cells and kinase activity underpinning the bending of developing cochlear duct.

We showed that the mode of luminal-basal nuclear migration switches at a position in the continuous pseudostratified epithelium (Fig. 2D’), which controls the curvature of the MEL (Fig. 2J-M). To our knowledge, we have provided the first experimental evidence that nuclei stall at the luminal side of the pseudostratified epithelium during IKNM in normal development (Norden, 2017). The luminal nuclear stalling results in luminal expansion, to some extent, because of physical space occupation even in the curved pseudostratified epithelium, while it remains unclear whether the luminal nuclear stalling results from the MEL curvature, i.e., the convexity of the luminal side. Therefore, we propose luminal expansion driven by luminal nuclear stalling as a mechanism for sculpting curves in pseudostratified epithelium, in addition to the actomyosin-based basal shrinkage that has been previously reported in zebrafish neuroepithelium (Sidhaye and Norden, 2017; Yanakieva et al., 2019). One important remaining question is what causes the space-dependent mode transition of nuclear movement in the continuous tissue. It has been previously shown that knock-down of cell-surface TAG-1 (transient axonal glycoprotein-1) impedes basalward nuclear movement during IKNM in the mouse ventricular zone, which ultimately leads to overcrowding of neural progenitor cells in the luminal side, and a severe cortical dysplasia(Okamoto et al., 2013). It has been proposed that cytoskeletal machinery, including actomyosin and microtubules, regulates basalward nuclear movement (Kosodo et al., 2011; Norden, 2017; Norden et al., 2009). This provides an important basis for the identification of the molecules responsible for the nuclear behavior in the MEL at the supra-cellular level, which needs to be clarified in the future. In addition, unknown factors providing these molecules with positional information should be explored to understand the mode transition of nuclear movement at the tissue level.

In the present study, the establishment of long-term imaging techniques and biosensors for protein kinase activity has led to the discovery of unexpected spatiotemporal patterns of cell movement and ERK activity in the developing cochlear duct. Previously, we and others have observed intercellular ERK activation waves in the epithelium, such as migrating Madin-Darby canine kidney (MDCK) cells (Aoki et al., 2017; Hino et al., 2020), developing drosophila tracheal placode (Ogura et al., 2018), and wounded murine skin (Hiratsuka et al., 2015). We have also proposed an ERK-mediated mechanochemical feedback system, in which cell extension activates ERK followed by ERK-triggered cell contraction (Boocock et al., 2020; Hino et al., 2020), which can explain the symphony between cell movement and ERK activity.

The ERK activation wave speed in the developing cochlear duct was 0.42 μm min^−1^ (Fig. 4F), which is significantly slower than in MDCK cells and wounded mouse epidermis, where it proceeds at 2.5 μm min^−1^ and 1.4 μm min^−1^, respectively (Aoki et al., 2017; Hino et al., 2020; Hiratsuka et al., 2015). Interestingly, when normalized to the cell lengths of the developing cochlear duct (4 μm), MDCK cells (20 μm), and the basal cells of the mouse epidermis (10 μm), the wave speed of the developing murine cochlear duct, which is 6 cells hr^−1^, is comparable with that of MDCK cells, which is 7 cells hr^−1^, and that of wounded adult murine skin, which is 8 cells hr^−1^. Our findings couple multicellular flow and ERK activation waves in the cochlea, corroborating the findings of earlier studies (Boocock et al., 2020; Hino et al., 2020), and supporting the existence of a general regulatory mechanism for collective cell migration during tissue morphogenesis.

Our 3D time-lapse imaging revealed coherent helical cell flow from the base-floor to the apex-roof of the lateral side of the cochlear duct. Cell flow analysis revealed that the rate of base-to-apex cell flow (0.24 μm min^−1^, Fig. S3E) exceeds that of the duct elongation speed (0.13 μm min^−1^, Mov. 5). Thus, the cell flow rate may be sufficient to compensate for the lateral tissue growth. We speculate that this ERK-mediated cell advection originating from the heterogeneity of cell proliferation causes consistent mediolateral differential growth at the tissue scale, and results in cochlear bending. In support of this, knockout of *Shh* causes a significant decrease in the number of proliferating cells at the base of the cochlear duct, and causes the shortening of the cochlear duct (Bok et al., 2013). As SHH is secreted mainly from the spiral ganglions located in the central axis of the cochlea (Bok et al., 2013), further investigations on the interaction between the cochlear duct and these ganglions will provide a better understanding of cochlear morphogenesis.

Overall, we visualized multicellular behavior underlying the bending of the cochlear duct during development using deep tissue live imaging. This contributes to a better understanding of symmetry breaking in tissue morphogenesis during development and in generation of inner ear organoids (Koehler et al., 2017, 2013). The live imaging technique used in the present study forms the basis for further analysis of the interplay between morphogenesis and cell fate decisions during cochlear development (Cohen et al., 2019; Tateya et al., 2019).

## Acknowledgements

This work was supported by the JSPS KAKENHI 17KT0107 and 19H00993, by the JST PRESTO JPMJPR1949 and CREST JPMJCR1654, and by the Medical Research Support Center of Kyoto University. We would like to thank Akane Kusumi for technical assistance, and Edouard Hannezo, Naoya Hino, and Yoshiko Takahashi for fruitful discussion.

## Author contributions

Conceptualization, T.T., T.H.; Methodology, M.I., T.T.. T.H.; Software, M.I., T.H.; Validation, M.I., T.H.; Formal analysis, M.I., T.H.; Investigation, M.I., T.H.; Resources, M.I., M.M., T.H.; Data curation, M.I., T.H.; Writing - original draft, M.I., T.H.; Writing - review & editing, T.T., M.M., T.H.; Visualization, T.H.; Supervision, M.M., T.H.; Project administration, T.H.; Funding acquisition, M.M., T.H.

## Materials and Methods

### (1) Experiments

#### Animals

For FRET imaging, we used the transgenic mice that ubiquitously express an ERK biosensor with a long flexible linker (hyBRET-ERK-NLS) reported elsewhere(Harvey et al., 2008; Komatsu et al., 2018, 2011). For simultaneous imaging of F-actin and nuclei, we crossed Lifeact-EGFP (Riedl et al., 2010) and R26-H2B-mCherry(Abe et al., 2011). Lifeact-EGFP mice were generously provided by Takashi Hiiragi from EMBL Heidelberg and R26-H2B-mCherry mice were provided from RIKEN Large (CDB0204K). Otherwise, we used ICR mice purchased from Japan SLC, Inc. We designated the midnight preceding the plug as embryonic day 0.0 (E0.0), and all mice were sacrificed by cervical dislocation to minimize suffering. All the animal experiments were approved by the local ethical committee for animal experimentation (MedKyo 19090 and 20081) and were performed in compliance with the guide for the care and use of laboratory animals at Kyoto University.

#### Antibodies

The following primary and secondary antibodies were used for immunofluorescence: anti-E-cadherin rat antibody (Cell Signaling Technology, #3195, 1:100 dilution), anti-Myosin-VIIa rabbit polyclonal antibody (Proteus BioSciences Inc., #25-6790, 1:200 dilution), anti-Histone H3 (phospho S28) rat polyclonal antibody (Abcam, #ab10543, 1:200 dilution), Alexa Fluor 546-conjugated goat anti-rat IgG (H+L) antibody (Thermo Fisher Scientific, #A11081, 1:1000 dilution), Alexa Fluor 647-conjugated goat anti-rabbit IgG (H+L) antibody (Abcam, #ab150079, or Thermo Fisher Scientific, #A21247, 1:1000 dilution).

#### Small-molecule inhibitors

The following chemicals were used: Mitomycin C (Nacalai Tesque, #20898-21) and PD0325901 (FUJIFILM Wako Pure Chemical Corporation, #162-25291).

#### Whole-tissue staining and imaging

The cochleae were gently freed from the capsule and the staining and clearing were performed according to an earlier study (Hirashima and Adachi, 2015). Briefly, the samples were fixed with 4% PFA in PBS overnight at 4°C and then blocked by incubation in 10% normal goat serum (Abcam, #ab156046) diluted in 0.1% Triton X-100/PBS (PBT) for 3 h at 37°C. The samples were treated with primary antibodies overnight at 4°C, washed in 0.1% PBT, and subsequently treated with secondary antibodies conjugated to either Alexa Fluor 546 or Alexa Fluor 647 overnight at 4°C. For counter staining of nucleus, we mixed Hoechst 33342 (5 μg/ml, Dojindo Molecular Technologies, #H342-10) or DAPI (Dojindo Molecular Technologies, #D523-10, 1:200 dilution). The samples were mounted with 10 μL of 1% agarose gel onto a glass-based dish (Greiner Bio-One, #627871) for stable imaging. Then, the samples were immersed with the BABB (benzyl-alcohol and benzyl-benzoate, 1:2, #04520-96 and # 04601-65, Nacalai Tesque) solution or CUBIC-R+ (Tokyo Chemical Industry Co., # T3741) solution for optical clearing. Images were acquired using the confocal laser scanning platform Leica TCS SP8 equipped with the hybrid detector Leica HyD with the ×40 objective lens (NA = 1.3, WD = 240 μm, HC PL APO CS2, Leica) and the Olympus FluoView FV1000 with the ×30 objective lens (NA = 1.05, WD = 0.8 mm, UPLSAPO30XS, Olympus).

#### EdU assay

For EdU incorporation to embryos, 200 μL of 5 mg/mL EdU in PBS was intraperitoneally injected to pregnant mice 30 min prior to dissection. For the incorporation to dissected cochleae, 10 μM of EdU was treated into the samples 1 hour prior to the chemical fixation. Before EdU detection, whole-tissue immunofluorescence of E-cadherin and counter nuclei staining with DAPI were performed. EdU was detected using the Click-iT® EdU Imaging Kits (Thermo Fisher Scientific, #C10340). The samples were optically cleared with CUBIC-R+ and images were acquired by confocal microscopy as described above.

#### Mosaic cell labeling

DNA solutions of the pCAG-GFP vectors (0.7 μg/μL) in PBS with 0.1% Fast Green FCF (Sigma-Aldrich, #F7252-5G) were injected into the lumen of an inner ear dissected from E13.5 embryos through a fine glass capillary tube under a stereo microscope (SZ61, Olympus). After injection of DNA solution, the cochlea was sandwiched by a pair of tweezer type electrodes (Nepa Gene Co., #CUY650P5) and the DNA was electrotransferred using the NEPA21 Super Electroporator (Nepa Gene Co.). Three 5 msec square pulses (175 V, +) with a 10% decay rate at intervals of 50 msec were applied as poring pulses, followed by three 50 msec square pulses (10 V, +/−) with a 40% decay rate at intervals of 50 msec as transfer pulses.

#### Surgical separation

We dissected cochleae from embryos at E17.5 and manually separated a cochlear duct into the medial and lateral side. The images in Movie 3 were acquired using the stereomicroscope (SZX16, Olympus) with a cooled color CCD camera (DP73, Olympus).

#### Explant cultures

We cultured the dissected cochleae without removing the capsule otherwise noted. The cochleae were mounted on a the 35 mm glass based dish (Iwaki, #3910-035) with 1 μL of growth factor reduced Matrigel (Corning, #356231), and filled with 2 mL of a culture medium including FluoroBrite DMEM Media (Thermo Fischer Scientific, #A1896701) with 1% GlutaMAX (Thermo Fischer Scientific, #35050061) and 1% N2 Supplement with Transferrin (Holo)(FUJIFILM Wako Pure Chemical Corporation, #141-08941). The samples were incubated at 37°C under 5% CO_2_.

#### Live imaging for explants

We used two different sample preparations for live imaging under ex vivo culture condition. For imaging of MEL, we completely removed the capsule and put the uncovered cochlear duct attached with spiral ganglion onto the glass-bottom dish whose apex side faces the glass. For long-term organ-scale imaging, we partially cut off the capsule adjacent to the apex tip of cochlear duct using tweezers carefully, and the semicircular canals were removed. The isolated cochlea was put onto the dish as described above. The former method allows us to take images MEL at single cell resolution but failed to achieve elongation and bending of cochlear duct. In contrast, the latter method recapitulates the cochlear duct morphogenesis but was unable to detect fluorescence signals in the medial side. For microscopy, we used an incubator-integrated multiphoton fluorescence microscope system (LCV-MPE, Olympus) with a ×25 water-immersion lens (NA=1.05, WD=2 mm, XLPLN25XWMP2, Olympus) and an inverted microscope (FV1200MPE-IX83, Olympus) with a ×30 silicone-immersion lens (NA=1.05, WD=0.8 mm, UPLSAPO30XS, Olympus). The excitation wavelengths were set to 840 nm, 930 nm, and 1040 nm each for CFP, Lifeact-EGFP, and R26-H2B-mCherry (InSight DeepSee, Spectra-Physics). Imaging conditions for Lifeact-EGFP and R26-H2B-mCherry were as follows – scan size: 800×800 pixels, scan speed: 4.0 μsec/pixel, IR cut filter: RDM690 (Olympus), dichroic mirrors: DM505 and DM570 (Olympus), and emission filters: BA495-540 for EGFP and BA575-630 for mCherry (Olympus). Imaging conditions for the FRET biosensor were as follows – scan size: 800 × 800 pixels, scan speed: 10 μsec/pixel, IR cut filter: RDM690 (Olympus), dichroic mirrors: DM505 and DM570 (Olympus), and emission filters: BA460-500 for CFP and BA520-560 for FRET detection (Olympus).

### (2) Quantification and Analysis

#### FRET image analysis

Image processing for FRET measurement was described elsewhere (Aoki et al., 2013). Briefly, the median filter of 3×3 window was processed to remove shot noises, and background signal was subtracted each in FRET channel and CFP channel. Then, the ratio of FRET intensity to the CFP intensity was calculated by a custom-made MATLAB (MathWorks) script.

#### Measurement of layer curvature and thickness

For 2D measurement of curvature and thickness, we first performed whole-mount immunofluorescence of E-cadherin to visualize the cochlear epithelium and acquired z-stack images by confocal microscopy as described above. Next, we manually traced the apical and basal side of epithelial cells on the middle horizontal section of the roof-floor axis. The extracted epithelial layer was named as the medial/lateral epithelial layer according to the side based on a given apex tip point. Then, the curve of the epithelial layer was obtained by the iterative skeletonization, and discrete points (*x*_*i*_, *y*_*i*_) were sampled along the curves at regular intervals of 15 μm. Finally, fitting the discrete points with a cubic spline function, the function *S*_*i*_ at an interval [*x*_*i*_, *x*_*i*+1_] is denoted as

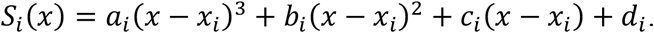

Due to a definition of curvature *κ*(*x*) = *S*′′(1 + *S*′^2^)^−3/2^, the curvature from the spline function was calculated as

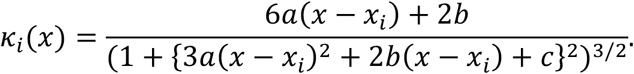

The curve of the convex/concave to the duct lumen was assigned as positive/negative in *κ*. We defined the layer thickness as a linear length connecting to the luminal and basal edge, which is vertical to the curve of the epithelial layer at sampling points.

#### Quantification of nuclear migration

We manually measured the center position of a daughter cell and its luminal and basal edge.

Based on the cross point where the luminal-basal line and center line of medial epithelial layer are intersected, we calculated the arc length from the duct tip to the nuclear position using a custom built MATLAB code.

#### Angle measurement of cell division orientation

First, we performed two-photon live imaging using FV1200MPE-IX83 as described above, and obtained volumetric images with a depth interval of 3 μm. In each mitotic event, we then manually measured position (x, y, z) of the two daughter cells 3 hours later after the mitosis in the fixed coordinate system O of the 3D image. Also, we manually acquired the position of the luminal and basal edge of the mother cell to define a position (x’, y’, z’) in a local coordinate system, O′, the origin of which is the middle point of the two daughter nuclei. Each orthogonal basis of the O′ coordinate was defined as follows: the apex-base axis in the MEL (x′), the surface normal of the MEL (y′), i.e., luminal-basal axis in the MEL, and roof-floor axis (z′) orthogonal to both x′ and y′ according to the right-handed system. With this local coordinate system, we finally calculated two angles φ and θ in the local sphere coordinate resulting from coordinate transformation from the O into the O′ system.

#### EdU intensity mapping

First, we separated 8-bit image stacks into two, roof and floor, based on z position at the middle point, and performed maximum intensity projection onto xy plane both for staining images for E-cadherin and EdU. Next, we binarized immunostaining signals for E-cadherin using Otsu’s method with morphological operations and detected periphery of the cochlear duct with the MATLAB function ‘bwperim’. With three reference points given manually, i.e., a) the duct tip, b) the end point of MEL and c) that of LEL, the MEL and LEL curve was each defined by connecting points between a) and b) and those between a) and c) along the duct periphery. Then, we marked points to make 20 bins at a constant distance each along the MEL curve and LEL curve. By connecting the marked point of MEL and that of LEL indexed by an order from the duct tip and dividing the lines into 10, we partitioned the cochlear duct into small regions for the measurement. Finally, we measured the averaged intensity of EdU signal within each region and normalized by 255.

#### Tissue flow and ERK activity

To calculate velocity fields of cells in cochleae, we performed particle image velocimetry (PIV)-based image processing using a free code MatPIV (a GNU public license software distributed by Prof. Kristian Sveen in University of Oslo) was applied to time-lapse images of the CFP channel. Velocity fields at time *T* was computed by displacement between *T* and *T*+*Δt*. *Δt* was set as the sampling rate, 12 min. The size of the interrogation window was set to 40 pixels, approximately 25 μm, corresponding to 4-5 cell diameter, and the window overlap was set to 50%. The obtained velocity data were then smoothened via median filtering to eliminate peaky noises. We then obtained the ‘tissue flow speed for the elongation’ from PIV velocity vector projected onto the apex-base line depicted in Figure 4D, and calculated the spatial derivative of the tissue flow speed for the elongation between two adjacent interrogation windows on the apex-base line according to the definition of a diagonal component of the strain rate. This quantity was smoothened using the MATLAB function ‘smooth’ to eliminate high frequency components and defined as the extension-shrinkage rate. As for the ERK activity, we set thresholds in CFP images using Otsu’s method within each interrogation window to extract cytoplasmic region and calculated the mean FRET/CFP ratio in the binarized region. The ERK activity rate was calculated as the time derivative of the ERK activity. Cross-correlation analysis was performed using the MATLAB function ‘xcorr’.

#### Statistical analysis

The number of cells or region of interests analyzed (n) and the number of biological replicates (N) are indicated in the figure legends. No particular statistical method was used to predetermine the sample size. A minimum of N=3 independent experiments were performed based on previous studies in the field. No inclusion/exclusion criteria were used and all analyzed samples were included in the analysis. No randomization was performed. Statistical tests, sample sizes, test statistics, and *P*-values were described in the main text. *P*-values of less than 0.05 were considered to be statistically significant in two-tailed tests, and were classified as 4 categories; * (*P*<0.05), ** (*P*<0.01), *** (*P*<0.001), and n.s. (not significant, i.e., *P* ≥ 0.05).

#### Software

For digital image processing, we used MATLAB (MathWorks) and Image J (National Institute of Health). For graphics, we used MATLAB (MathWorks), Imaris (Bitplane) and ImageJ (National Institute of Health). For statistical analysis, we used MATLAB (MathWorks).

### (3) Mathematical model

#### A-1. A mechanical model for MEL morphogenesis

We modeled multicellular dynamics within an epithelial layer using a vertex dynamics model (VDM) (Fletcher et al., 2014; Hirashima and Adachi, 2019; Nagai and Honda, 2001). Here we focused on two-dimensional section on apex-base and roof-floor axes as shown in Fig. 2A. In general, the 2D VDM model represents a single cell as a polygon, of which vertices are elementary points that constitute the cell shape, and a group of cells is regarded as a set of polygons shared by neighboring cells – we put four vertices, two of which are regarded as an luminal edge and the other two of which are as a basal edge.

In the VDM, the dynamics of position of vertex *i*, *r*_*i*_, obey the equation of motion based on the principle of least potential energy *U* as follows:

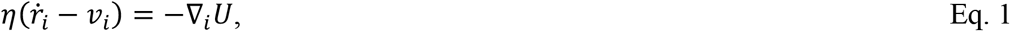

where *η* is a viscosity coefficient. *v*_*i*_ is a local velocity of vertex *i*, defined as 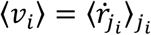, where *j*_*i*_ is an index of cells contacting to vertex *i*, 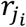 is a centroid of cell *j*_*i*_, and 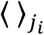 denotes averaging in *j*_*i*_ (Hirashima and Adachi, 2019; Mao et al., 2013; Okuda et al., 2014). For a potential energy as a minimum expression, we defined as

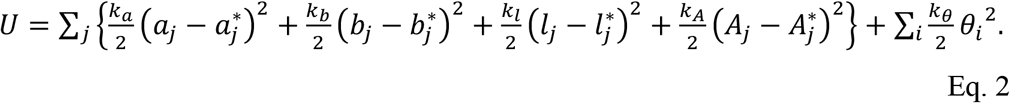

The first to third terms each represent a regulation of cell edge length in luminal/apical, basal, and lateral side with controlling parameters (*k*_*a*_, *k*_*b*_, *k*_*l*_), current length (*a*_*j*_, *b*_*j*_, *l*_*j*_, and the target length 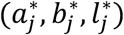. Target edge length of the luminal/apical and that of basal side are a function of nuclear position within the epithelial layer, described later. The fourth term represents cell area preservation with its coefficient *k*_*A*_, current cell area *A*_*j*_, and the target cell area 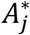. The fifth term represents bending energy of the luminal/apical side and that of basal side attributed to each vertex; *k*_*θ*_ denotes the bending rigidity and *θ*_*i*_ is an angle of luminal/apical edge or that of basal edge at a vertex *i*. Although the cell shape is relatively flexible in a pseudostratified epithelial tissue, this model framework is valid as several biological features can be incorporated, such as the nuclear position-dependent edge regulation and cell size preservation.

Each cell has a unique cell cycle, length of which *τ*_*div*_ is assumed to be equally assigned to all cells. We assume that a period of each cell cycle phase (G1, S, G2, and M) is partitioned as 11:8:4:1, and a cell cycle status or timer *τ*_*j*_, origin of which is defined at the end of cell division, i.e., a boundary of M-G1, is provided according to the distribution of cell cycle phase. We model that the cell timer *τ*_*j*_ in the cycle determines the distance from the luminal side of a cell, i.e., nuclear position *d*((0 ≤ *d*_*j*_ ≤ 1), with a parameter *γ* as follows:

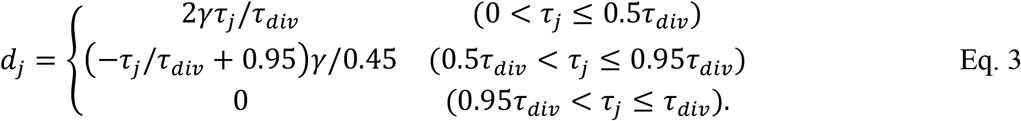

This means that nuclei move toward the basal side of cell during G1-S phase with a degree *γ* and get close to the luminal side before entering M phase. Of note, *γ* controls the degree of basalward movement as described in the main text. Once a cell divides, *τ*_*j*_ in one of daughter cells reset to zero and one in the other is set to a value stochastically chosen from 0 to 0.1*τ*_*div*_ with a uniform distribution to avoid perfect synchronization between neighboring cells.

Occupied cell area is dominant in the medial epithelial layer and the cell position along the luminal-basal axis should regulate the length of luminal edge and that of basal edge. Thus, we define the target length as the simplest linear function of the nuclear position with two parameters *ξ*_*max*_ and *ξ*_*min*_ as follows:

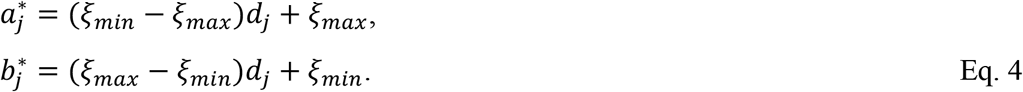

#### A-2. Numerical simulation

The ordinary differential equations were numerically solved by the forward Euler method with time step 0.01. The code was generated with MATLAB (MathWorks). Regarding initial condition, 20 rectangular cells, width/height of each which is 2.5/50 μm, are arrayed along a horizontal line. Standard parameter set is *η* = 1, *k*_*A*_ = 0.01, 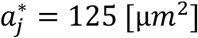, *k*_*a*_ = 1, *k*_*b*_ = 1, *k*_*l*_ = 1, *k*_*θ*_ = 30, 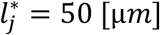, *τ*_*div*_ = 432, *γ* = 0.9, *ξ*_*max*_ = 5 [μ*m*], *ξ*_*min*_ = 1 [μ*m*], otherwise noted. 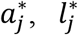, *γ*, *ξ*_*max*_, and *ξ*_*min*_ were determined from obtained images, and other parameters were determined empirically with some observations; at first, *η*, *k*_*a*_, *k*_*b*_, and *k*_*l*_ were set to 1 and then *k*_*A*_, *τ*_*div*_, and *k*_*θ*_ were determined to reproduce the MEL curvature observed in the curvature experimentally. Note that relative values rather than absolute ones are critical for the dynamics.

#### A-3. Detailed setting in virtual experiments for cell cycle arrest

For an investigation towards mimicking the developmental process, we introduced luminal stalling region, defined with an arclength along the epithelial layer *L* originated from the boundary of apex side. Assuming once the cells in the luminal stalling region undergo division, the daughter cells’ timer *τ*_*j*_ do not count up, meaning that those nuclei stay at the luminal side without reentering the cell cycle. This is due to experimental observations in which daughter cells do not undergo mitosis exclusively in the curved region within a short time scale, at least our observation window. For the cells in the non-luminal stalling region, *τ*_*div*_ was set stochastically chosen from a uniform distribution from 216 to 864 each after the cell division to incorporate variability and asynchronicity in nuclear dynamics. To recapitulate the mitomycin C treatment in simulation, the cell cycle was arrested if the cell timer *τ*_*j*_ was ranged in 0.25*τ*_*div*_ to 0.8*τ*_*div*_, corresponding to the mid G1 phase to the end of S phase, when the simulation time exceeded 700. For the control case, we evaluated the curvature of virtual MEL when the cell number reached at 100. For the case of mitomycin C treatment, we evaluated when the simulation time reached at 1300 because the averaged simulation time in the control case was 1298 (n=10).

#### B-1. Modeling oscillatory ERK activation waves and cell flows

We built a minimal 1D mechano-chemical coupling model for the collective cell migration based on our previous studies (Boocock et al., 2020; Hino et al., 2020). Cells, each indexed as *j*=1,…,*N*, are represented as a chain of springs, whose junctions including the boundaries are labeled as *i*=1,…,*N*+1, with elastic constant *k* and each cell generates contractile force at the rear side of the cell to move to the front with a force *F*. *i.e.*, the cell contractile force with *j*=*N* (*F*_*j*=*N*_) is regarded as the force at the junction *i*=*N* (*F*_*i*=*N*_). Because the epithelial cells adhere to neighboring cells and thus transmit the elastic force with viscous frictions ***η***_***c***_, the dynamics of cell collectives is represented as

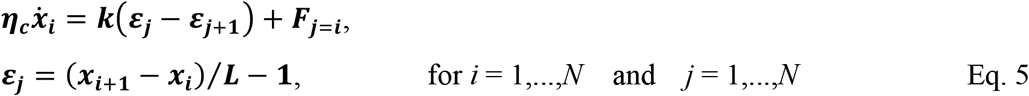

where ***ε*** is a cell strain and *L* is a typical cell length, i.e., 5 μm. At the front edge of the cells, *i.e.*, *i*=*N*+1, a self-propelling force *F*_*tip*_ is generated, reflecting an elongation of the duct tip. Since the cells respond to stretching as activating the ERK, a coupling between the cell kinematics and the ERK activity is formulated as

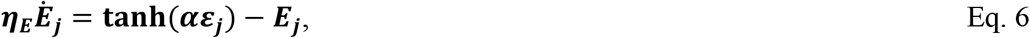

where *α* denotes a sensitivity parameter and ***η***_***E***_ denotes a timescale of the dynamics. Then, the ERK activity is converted to the self-contractile force represented as dynamics of the

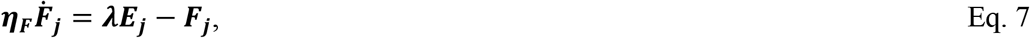

where ***λ*** denotes a controlling parameter of amplitude, and ***η***_***F***_ denotes a timescale.

As for the uncoupling regime, ERK activity was given as the following traveling waves instead of Eq. 6:

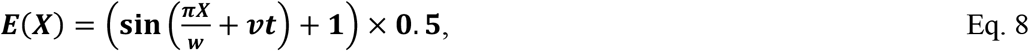

where *w* is the characteristic length of ERK activation and *v* is the ERK activation speed.

The parameters *w* was set as 84 μm, i.e., the wavelength of the ERK activity is 168 μm, from Figure 4E and *v* was set as 0.42 μm min^−1^ from Figure 4F.

#### B-2. Numerical simulation

The ordinary differential equations were numerically solved by the forward Euler method with time step 0.01 using the MATLAB. The number of cells *N* was set as 1000 and the one boundary *i*=1 was fixed and the other *i*=*N*+1 was the moving boundary condition. Biologically plausible parameter set was determined as ***η***_***c***_ = 40 [nN min μm^−1^], ***k*** = 20 [nN], *F*_*tip*_ = 6 [nN], ***α*** = 3, ***η***_***E***_ = 30 [min], ***η***_***F***_ = 10 [min], and ***λ*** = 9 [nN] according to the present study and a previous study (Serra-Picamal et al., 2012).

## Supplementary figure legends

**Figure S1 Morphological quantification of epithelial layer**

Curvature and thickness as a function of the arc length from the tip along the lateral epithelial layer (LEL, left) and the medial epithelial layer (MEL, right) at E13.5 to E15.5. Mean ± s.d. N=3.

**Figure S2 Luminal-basal nuclear movement in the MEL**

(A) Numerical investigation of the mathematical model. The curvature is plotted over ***γ***. Mean ± s.d. N=10. (B) Treatment of cochlea with 10 μM mitomycin C for 1 day. Maximum intensity projection of immunostained images for the anti-E-cadherin (white) and EdU labelling (magenta). Scale bar, 100 μm.

**Figure S3 ERK activity waves and cell flows**

(A) Roof plane (orange dotted line) in the cross section view of 3D ERK activity map in the cochlea at E12.5. Scale bar, 50 μm. (B) Time-lapse snapshots of ERK activity map in the roof plane. Time indicates the elapsed time of live imaging. Scale bar, 100 μm. (C) Time-lapse snapshots of tissue flow speed obtained by the PIV in the roof plane. The multicellular flows direct toward the elongation direction around the duct tip despite winding in the region away from the tip in the roof side. Scale bar, 100 μm. (D) Kymograph of ERK activity for the two samples. Horizontal axis indicates the position on the apex-base line and vertical axis indicates the elapsed time of live imaging. Dotted lines represent the oscillatory wave trains from the apex to the base. Scale bar, 100 μm. (E) Time-series data of ERK activity and the tissue flow speed for elongation in the three different ROIs. (F) ERK activity before and after the MEK inhibitor PD0325901 treatment at 1 μM. Scale bar, 20 μm.

**Figure S4 Mathematical model for ERK activation waves and cell flows**

(A, A’) Schematics for two regimes. ERK activity and cell deformation is reciprocally regulated in the mechano-chemical coupling regimes (A), but is unidirectionally regulated from the ERK activity to the cell deformation without closed feedbacks in the uncoupling regime (A’). (B, B’) Kymograph of the ERK activity. In the mechano-chemical coupling regime, the ERK activity propagation from the apex to the base can be generated (B), even without given ERK wave as shown in the uncoupling regime (B’). Scale bar, 100 μm. (C, C’) Time-series of ERK activity rate and extension-shrinkage rate in the coupling (C) and the uncoupling regime (C’). (D, D’) Cross-correlation between the extension-shrinkage rate and the ERK activity rate in the coupling (D) and the uncoupling regime (D’). The lag time in (D) is −28 min, compatible with the experiments, while the lag time in (D’) is −2 min, almost no lag time.

## Movie legends

**Movie S1 Real-time movie showing surgical separation of the cochlear duct at E17.5, related to Figure 1.**

After separation, the medial and lateral tissue are placed on the left and the right, respectively. Note that the medial tissue remains curled even after being straightened.

**Movie S2 Time-lapse movie of MEL in the cochlear duct at E14.5, related to Figure 2.**

Representative nuclei are labelled when those reach at the luminal surface. Green and magenta each denote the nucleus in the flat region and in the curved region. Time interval is 5 min.

**Movie S3 Model simulation for cell cycle arrest, related to Figure 2.**

Color in the epithelial layer indicates the curvature and that in the circles indicate the cell cycle state. Interkinetic nuclear migration is stopped at a timing of cell cycle arrest shown in the right panel.

**Movie S4 3D ERK activity map of the cochlear at E12.5, related to Figure 4.**

Color indicates the ERK activity level shown in Figure 4A. Views from roof side at initial, and ones from apex side at last.

**Movie S5 Time-lapse movie of the ERK activity and multicellular tissue flow, related to Figure 4.**

Color in the left panel indicates the ERK activity level shown in Figure 4B and that in the right panel indicates the PIV speed shown in Figure 4C. Views at the floor plane indicated in Figure 4A’. Time intervals is 12 min.

**Movie S6 3D time-lapse imaging of ERK activity in the lateral side of the cochlear duct at E12.5, related to Figure 4.**

Color indicate the ERK activity level shown in Figure 4J. Views from the lower left corner of Figure 4B. Green arrow indicates the ERK activation peak.

