## Supplementary figures and images for "Interplay between medial nuclear stalling and lateral cellular flow underlies cochlear duct spiral morphogenesis"

### Supplementary Figure

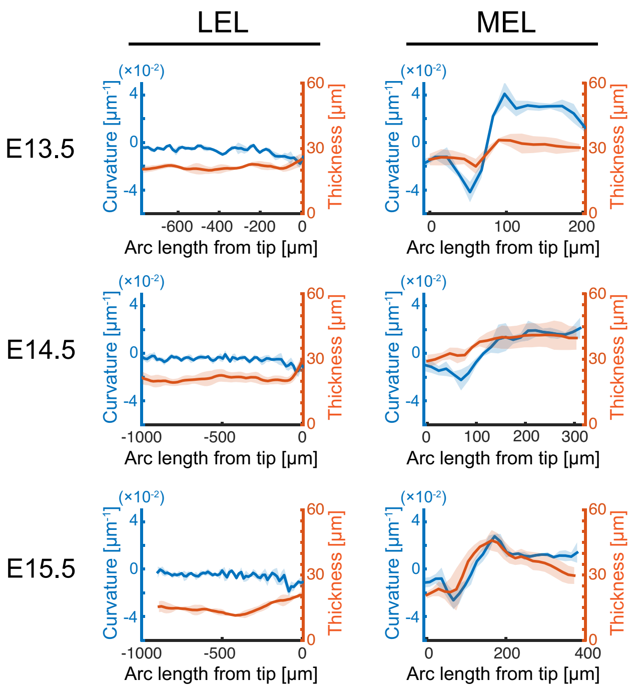

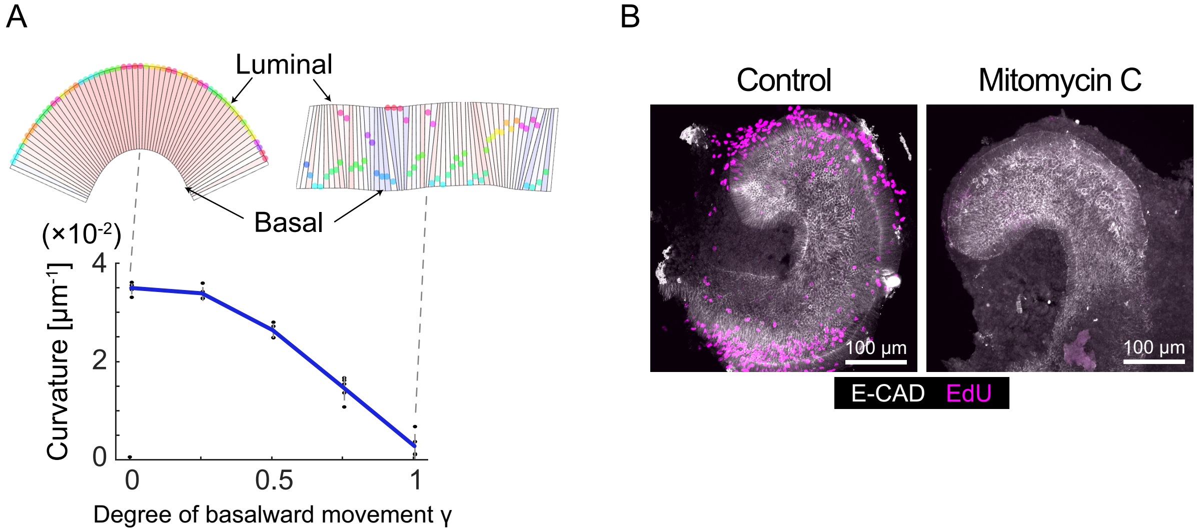

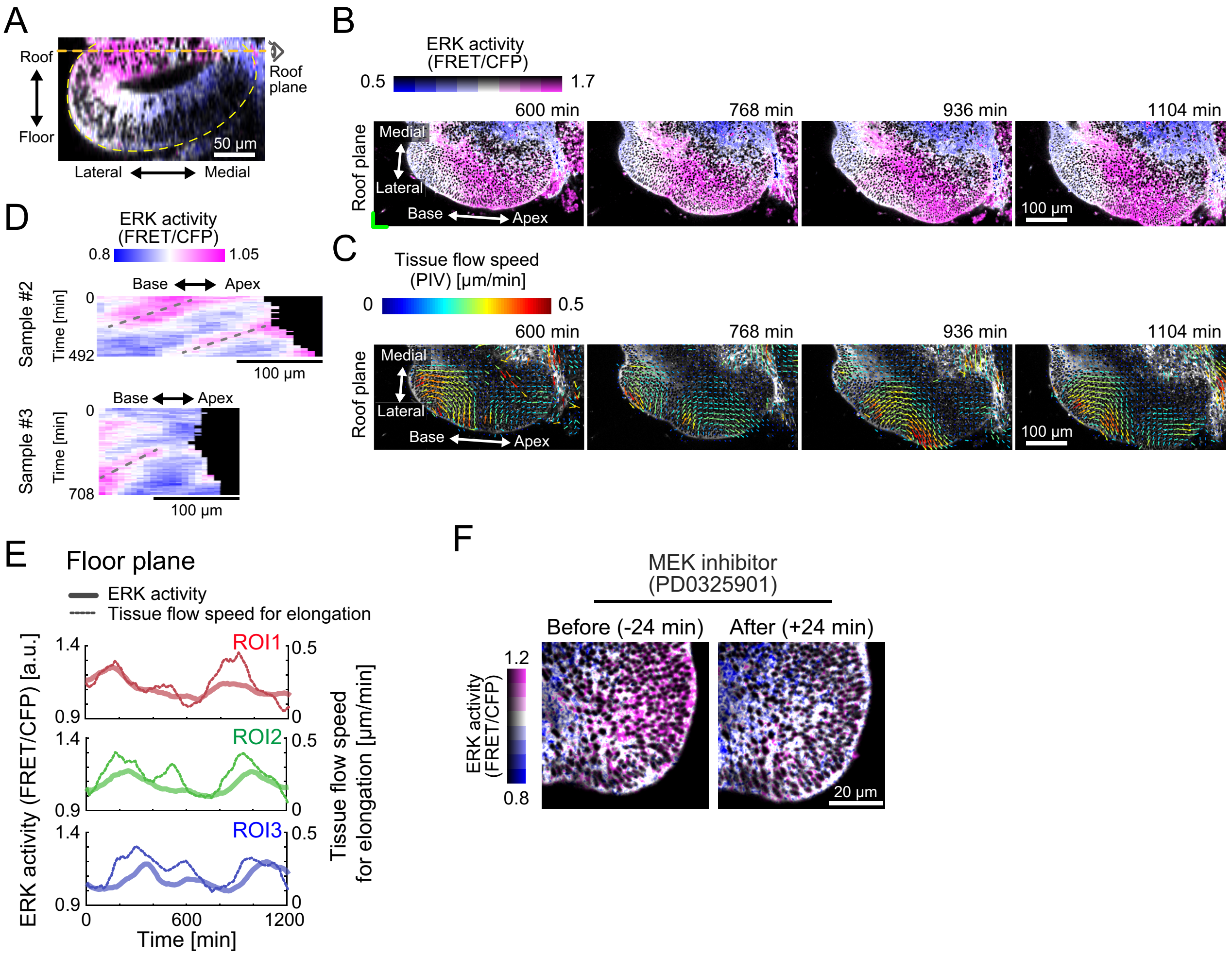

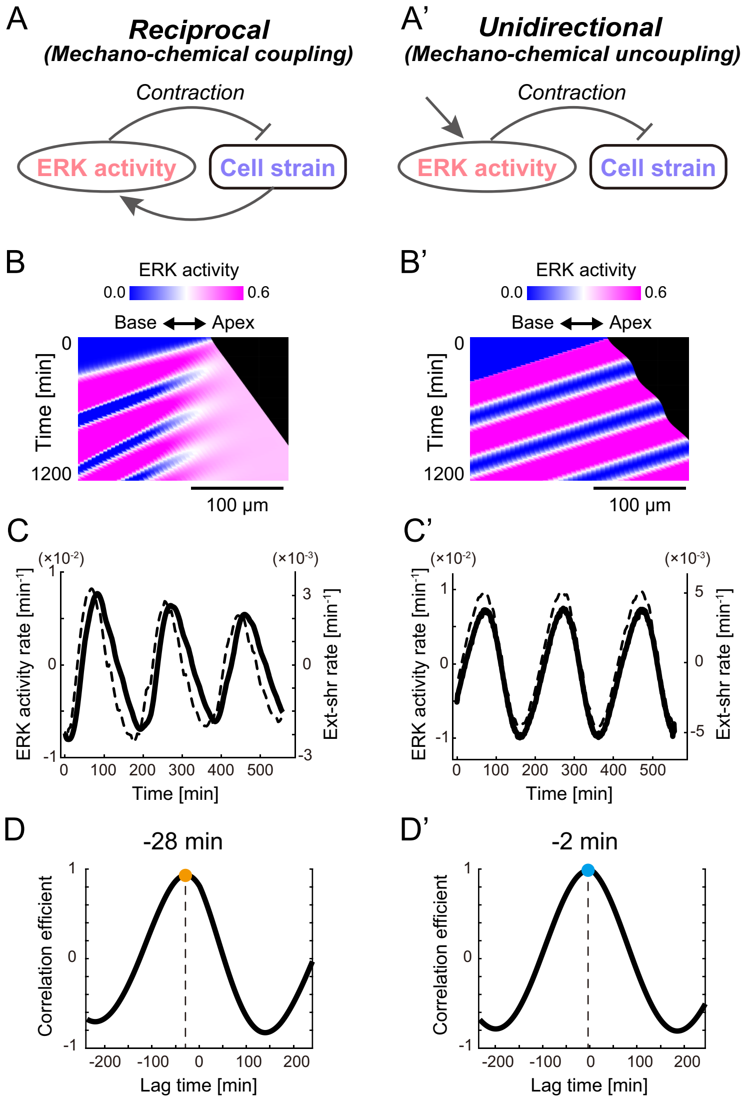
